# First description of two immune escape indian B.1.1.420 and B.1.617.1 SARS-CoV2 variants in France

**DOI:** 10.1101/2021.05.12.443357

**Authors:** Vincent Garcia, Véronique Vig, Laurent Peillard, Alaa Ramdani, Sofiane Mohamed, Philippe Halfon

**Affiliations:** Laboratoire Alphabio, Marseille; Service de Médecine Interne et de Maladies Infectieuses, Hôpital Européen, Marseille; Agence Régionale de Santé, Marseille, France; France ABL, Marseille, France

**Keywords:** N440K, E484Q/L452R SARS-CoV2, High throughput sequencing, Indian variant

## Abstract

Following the outbreak of the SARS-CoV2 virus worldwide in 2019, the rapid widespread overtime of variants suggests today an undergoing positive selection of variants which could potentially provide advantageous genetic property of the virus. Numerous variants have already been described across different countries including N501Y, E484K or L452R mutations on gene coding to spike protein. Most recently, 2 new Indian variants with N440K and E484Q and L452R mutations associated with impaired antibody response and immune reactions were identified in India. The potential consequences of emerging variants are increased transmissibility, increased pathogenicity and the ability to escape natural or vaccine-induced immunity.

We described for the first time in France both variants: the N440K immune escape variant within a new strain detected in France in a couple of patients who did not have any history of travel abroad and the new E484Q and L452R Indian variant from a patient travelling from Indian to Marseille to embark on a ship as a crew member.

Such study of the circulating viral strains and their variants within the increasing number of infected people worldwide will provide further insights into the viral dissemination. Hence, real time close monitoring variant could help the scientific community to prevent fast-spreading and raise alarms towards potentially harmful variants.

## Introduction

Since the beginning of the SARS-CoV2 virus pandemia, recent studies exploring the role of novel coronavirus genetic variations in escaping immune response have shed light on the possible mechanisms of the pathogen to evade antibody response and immune reactions (1,2). The best way to control the potential damage is to monitor the prevalence of circulating variants and the emergence of new mutations throughout extensive viral genome sequencing within the infected population. Such surveillance will then allow to prevent the spread of new variants potentially more infectious and harmful. Indeed, the use of high throughput sequencing allows the identification of emerging variants responsible for epidemic rebound or localized clusters of infected patients conventional RT-PCR assay would not identify that.

Beside the main variants previously described N501Y, E484K or L452R mutations on gene coding to spike protein (3,4), in several countries, 2 new variants with N440K and E484Q and L452R mutations associated with impaired antibody response and immune reactions were identified in India 5,6.

In this context, Alphabio’s medical laboratory in Marseille (France) started a real time genomic survey of the Sars-CoV-2 circulating variants based on high-throughput whole genome sequencing in collaboration with the Agence Régionale de Santé PACA, in Marseille. This collaboration aims to characterize specific clusters and identify new emerging variants responsible for increased infectivity and/or immune escape. Here, we describe for the first time, the characterization of the two immune escape variants: N440K detected in a couple of patients and the E484Q and L452R Indian variant from a patient travelling from Indian to Marseille to embark on a ship as a crew member in France.

## Materials and Methods

From the first of March 2021, each new SARS-CoV2 samples sequenced at Alphabio by RT-PCR were selected on the basis of the following criteria: i) the first positive sample for each patient, ii) threshold for SARS-CoV2 sample was set at a CT < 28, iii) Samples with RT PCR variant screening results either suspected of being 20H/501Y.V2 - 20J/501Y.V3 or negative for 20I/501Y.V1, 20H/501Y.V2 and 20J/501Y.V3.

Clinical samples from nasopharyngeal swabs were extracted using the Chemagic 360 (Perkinelmer Inc., USA). Detection of the SARS-CoV-2 was performed using the CE-IVD UltraGene Combo2screen rt-PCR SARS-CoV-2 (E/N) assay (ABL, France). Screening of the three main variants 20I/501Y.V1, 20H/501Y.V2, 20J/501Y.V3 was performed using the VIROBOAR SPIKE assay (Eurofins Genomics, Germany). SARS-CoV-2 whole genome was amplified on selected samples using the DeepChek^®^ assay Whole Genome SARS-CoV-2 Genotyping V1 (ABL, France) and sequenced using the DeepChek^®^ NGS Library preparation V1 (ABL, France) as per the manufacturer’s protocol. The resulting libraries were sequenced using the MiSeq system (Illumina, USA) and the run was performed generating 2 × 251 bp read length data during a 39-h run time. Filtered reads were mapped against SARS-CoV-2 reference using DeepChek^®^ - Whole genome SARS-CoV-2 algorithms (ABL, Luxembourg). To generate alignment statistics, coverage data and consensus sequence MicrobioChek software platforms (ABL, Luxembourg) were used. A phylogenetic tree was constructed using NextClade V. 0.14.2 webservice.

## Results

The first N440K escape variant was isolated from a 36-year-old French woman and her husband, 37-year-old that were tested positive for the first time for SARS-CoV2 with a cycle threshold at 16 on the 16th of March, 2021 at Alphabio’s medical biology laboratory, Hôpital Européen, Marseille. Both had mild symptoms including asthenia and cough without comorbidity or severity’s factors and were recovering. None of them was vaccinated against the COVID-19 virus and they did not have any history of travel abroad.

The second E484Q - L452R variant was isolated from a 37 –year-old Indian man from Mumbai arrived in Marseille on 16th of March, 2021 to embark on a ship as a crew member. On the 19th of March, 2021, he had mild symptoms including asthenia and cough without comorbidity or severity’s factors. He was tested positive for the first time for SARS-CoV2 with a cycle threshold at 15 and was immediately isolated in quarantine in a city hotel isolated.

A RT PCR of variants screening was performed and results showed that both strains did not belong to 20I/501Y.V1, 20H/501Y.V2 or 20J/501Y.V3 variants. Miseq sequencing generated a total of 7.2 Gb with 88 % cluster passing filter and 76% quality cutoff of (QC) 30.

Analysis of whole genome sequencing of the first one revealed 14 missense mutations including a T22882G (S: N440K) variant, as illustrated in Figure 1A. PANGO lineage assigned for this genome was **B.1.1.420**. The genomic sequence has been deposited in GISAID (assignation: 1623852).

**Figure 1.**
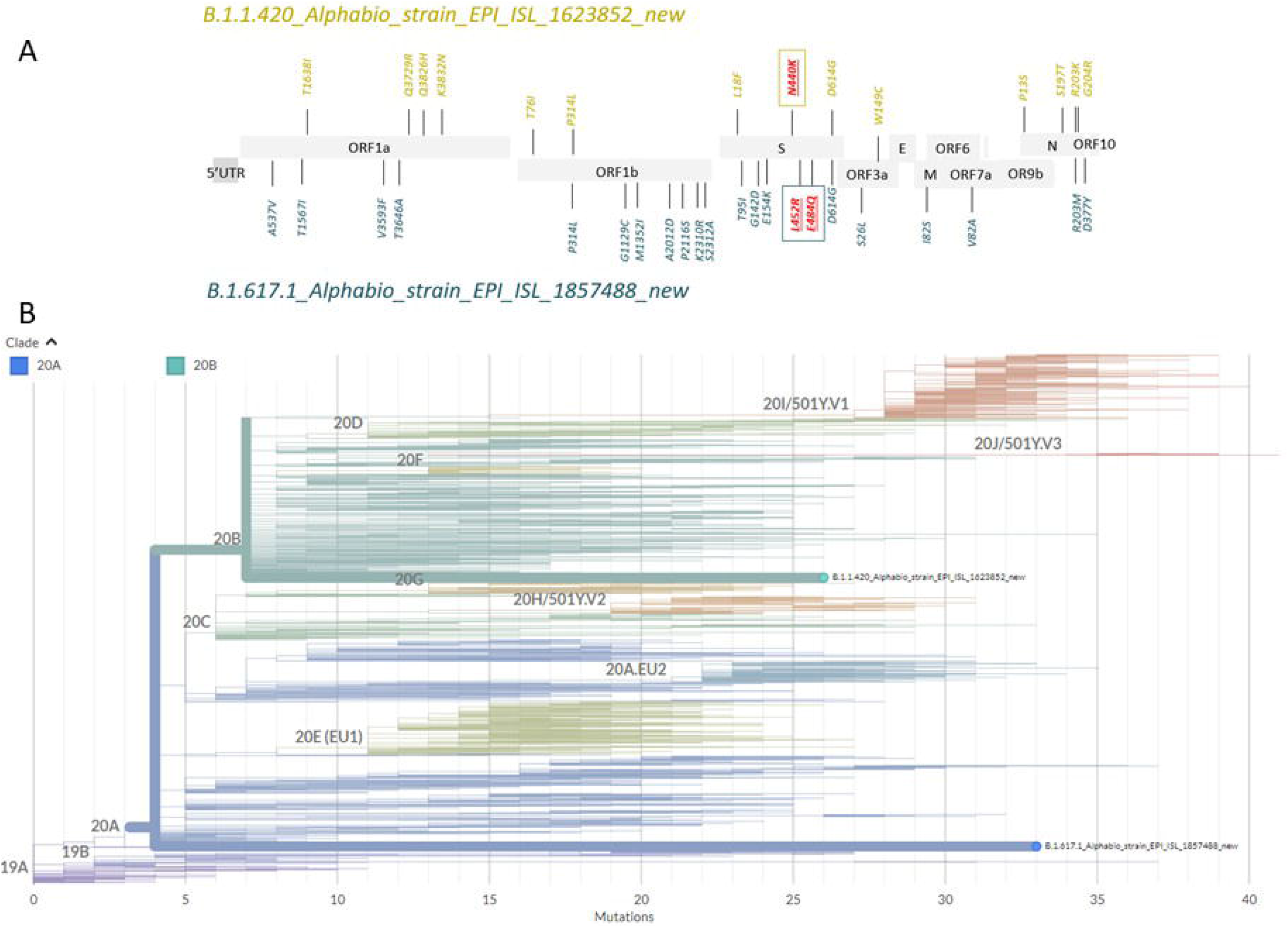
**A:** Genetic mutations in the genome isolate of SARS-CoV-2 infections. **B.** Phylogenetic tree including our N440K strain among different SARS-COV2 clades.

Analysis of whole genome sequencing of the second one revealed 22 missense mutations including a G23012C (S:E484Q) and T22917G (S:L452R), as illustrated in Figure 1A. PANGO lineage assigned for this genome was **B.1.617.1**. The genomic sequence has been deposited in GISAID (assignation: 1623852). Phylogenetic analysis showed that these strains belongs to Clade 20B and 20A respectively and is distant from other variants on concern known (Figure 1B).

## Discussion

Accurate and timely detection of new variants that may show greater infectivity or worse clinical symptoms, including immune escape, will be extremely important to prevent a worsening of the pandemia. In this study, the discovery of this first N440K variation isolated in France within infected subjects presenting no history of recent travelling highlights the importance of monitoring the impact of new viral variants. This N440K mutation, previously described in India, has been reported to be resistant to class 3 monoclonal antibodies (mAbs) C135 and REGN10987 that are candidates for clinical development (5). Both C135 and REGN10987 mAbs have been shown to have interactions focused on the N440 residue of the Spike protein and the proximity of the N440 residue to the structural epitope of the mAbs potentially confers loss of binding and resistance to the neutralizing effect of the mAbs. This variant is also reported to have an enhanced binding affinity to the ACE2 receptor in humans and may have the potential of a higher transmission rate. This new **B.1.1.420** genome sequence was deposited on the global database, GISAID. Studies looking at the N440K variant have not yet established weither or not the mutation affect the replicative fitness of the virus. However, as a new emerging variant, its low prevalence could also be explained by the lack of sequencing in the infected population carrying this mutation thus leading to an underestimation of the spread of this variant. Prior studies report that SARS-CoV-2 is rapidly spreading with numerous variants scattered across the globe. Given the high mutation rate of RNA viruses, it could be hypothesized that the emergence of the N440K variant is the result of an intrahost evolution.

The identification of the **B.1.617.**1 genome with the combination of the two RBD mutations E484Q or L452R noted in this study could affect the neutralization of the select mAbs (5). This case also raises the question of the lack of real control over transfers of people from areas where variants of concern are circulating. This patient had been tested before boarding and was able to transit unchecked to France where checks as many travelers as possible but without being exhaustive.

The rapid increase of infected people will provide more genome samples and could soon offer further insights into the mechanisms behind the viral dissemination of the SARS-CoV-2 worldwide. Hence, more coronavirus genomes need to be sequenced across the globe to accurately identify the emergence of these mutants and other variants. Considering the immune escape conferred by the N440K, E484Q and L452R mutation, these new variants must be further surveyed to avoid fast-spreading and raises alerts if it was considered to be soon a variant contributing to accelerate the spread of the virus in human populations.

## Ethical Issue

Informed consents were obtained from patients with agreement to use their data for this case report, in accordance with the Helsinki Declaration

## Conflict of interest and financial disclosure

The opinions expressed by authors contributing to this journal do not necessarily reflect the opinions of the Centers for Disease Control and Prevention or the institutions with which the authors are affiliated.

## Funding

ARS PACA Funding

## Notes

### Competing Interest Statement

The authors have declared no competing interest.

